# Standard selection treatments with sulfadiazine limit *Plasmodium yoelii* host-to-vector transmission

**DOI:** 10.1101/2022.01.13.476179

**Authors:** Kelly T. Rios, Taylor M. Dickson, Scott E. Lindner

**Author notes:** Corresponding Author. Scott E. Lindner.

## Abstract

Some early antimalarial drugs have been repurposed for experimental applications, thus extending their utility well beyond the point when resistance becomes prevalent in circulating parasite populations. One such drug is sulfadiazine, which is an analog of p-aminobenzoic acid (pABA), and acts as a competitive inhibitor of dihydropteroate synthase, which is an essential enzyme in the parasite’s folate synthesis pathway that is required for DNA synthesis. Sulfadiazine treatment of mice infected with *P. yoelii* and *P. berghei* is routinely used to enrich for gametocytes by killing asexual blood stage parasites, but it is not well known if the exposed gametocytes are perturbed or if there is a detrimental effect on transmission. To determine if there was a significant effect of sulfadiazine exposure upon host-to-vector transmission, we transmitted *Plasmodium yoelii* (17XNL strain) parasites to *Anopheles stephensi* mosquitoes and evaluated the prevalence of infection (percent of mosquitoes infected) and intensity of infection (number of oocysts per infected mosquito) under different sulfadiazine treatment conditions of the mouse or of the mosquitoes. We observed that parasites exposed to sulfadiazine either in the mouse host or in the mosquito vector had a reduction in both the number of mosquitoes that became infected and in the intensity of infection compared to untreated controls. We also observed that provision of freshly prepared pABA in the mosquito sugar water could only marginally overcome the defects caused by sulfadiazine treatment. In contrast, we determined that gametocytes exposed to sulfadiazine were able to be fertilized and develop into morphologically mature ookinetes *in vitro*, and thus that sulfadiazine exposure in the host may be reversible if the drug is washed out and the parasites are supplemented with pABA in the culture media. Overall, this indicates that sulfadiazine dampens host-to-vector transmission, and that this inhibition can only be partially overcome by exposure to fresh pABA *in vivo* and *in vitro*. Because gametocytes are of great interest for developing transmission blocking interventions, we recommend that less disruptive approaches for gametocyte enrichment be used in order to study minimally perturbed parasites.

## Introduction

Sulfa drugs have been used to treat human-infectious *Plasmodium falciparum;* sulfadoxine, in combination with another antifolate pyrimethamine, had been used extensively as an antimalarial therapy in endemic areas [1]. The widespread emergence of resistant parasites after drug pressure in clinical samples and in *in vitro* cultures makes these antifolates inappropriate for antimalarial monotherapies [2–4]. Because of this, sulfadoxine-pyrimethamine is now provided with artesunate as a WHO-recommended first-line combination therapy for the treatment of *P. falciparum* malaria in the WHO SE Asia regions [5].

Though sulfa drugs may not be as effective for treating human malaria infections today, sulfadiazine has been adapted as a commonly used tool to study rodent-infectious malaria parasites as it selectively kills the actively replicating asexual blood stages of the parasite and effectively enriches for sexual stage gametocytes [6]. Sulfa drugs act as antifolates by competitively inhibiting the interaction of an essential enzyme in the *de novo* folate synthesis pathway, dihydropteroate synthase (DHPS), with its substrate p-aminobenzoic acid (pABA). Antifolate drugs are effective against *Plasmodium* as they cannot use preformed folates like their hosts can and thus require *de novo* folate synthesis for the downstream generation of nucleic acids for DNA replication [7–8]. As such, actively replicating parasites, like those in asexual blood stages, are killed by sulfadiazine exposure, while non-replicating gametocytes survive and may be enriched by this treatment. *Plasmodium* parasites mainly source pABA from their hosts, though *de novo* pABA synthesis in *P. berghei* was recently observed when pABA in the rodent host diet was restricted [9]. In agreement with this, earlier work reported that pABA-deficient diets in rodent hosts is responsible for poor parasite growth and infection, indicating that pABA is an essential host-derived nutrient [10–11]. Indeed, newborn mice on naturally pABA-deficient milk diets suppressed parasitemia with Py17XNL infection and removal of pABA from the rodent diet reduced parasite load [12].

Therefore, it is notable that pABA is present in normal laboratory mouse feed at levels that allow for asexual blood stages to progress without additional supplementation (~175 ug/kg in conventional mouse feed) [9]. Similarly, laboratory-reared mosquitoes are commonly supplemented with pABA (0.05% w/v) in their sugar water to enhance oocyst numbers in transmission experiments [13].

Commonly, treatment with sulfadiazine to enrich for *P. berghei* or *P. yoelii* sexual blood stage parasites is accomplished by providing 10-30 mg/L (30-120 μM) sulfadiazine in the rodent host’s drinking water for 24-48 hours prior to parasite collection [14]. Despite this, little is documented regarding the possible downstream effects of sulfadiazine treatment on gametocytes, their host-to-vector transmission, and the early events of mosquito stage development. It is feasible that these parasite stages could be impacted by sulfadiazine, as they are known to undergo DNA replication in early mosquito development during gametogenesis, zygote-to-ookinete maturation, and in later development during sporogony [15]. *Plasmodium* transmission studies starting from the 1940s have indicated that there may be an effect of sulfadiazine exposure on host-to-vector transmission, though this early work used a variety of mosquito vectors and *Plasmodium* species combinations and was limited in the number of mosquitoes tested and their analyses of transmission [13, 16–20]. Perhaps because of this, these studies showed some inconsistencies. For example, work on *P. gallinaceum* transmission to *Aedes aegypti* showed that sulfadiazine inhibited sporozoite development [16–17] but follow up work a few years later suggested that sulfadiazine or sulfanilamide in the *Aedes aegypti* diet can even increase the insect’s susceptibility to *P. gallinaceum* oocyst development [18]. Work around the same time implied that *P. gallinaceum* oocyst growth in *Anopheles quadrimaculatus* is inhibited by sulfadiazine [19], but *P. vivax* transmission to *Anopheles stephensi* was not found to be inhibited by sulfadiazine [20]. Finally, work on *P. berghei* NK65 strain parasites in *Anopheles stephensi* showed that sulfadoxine exposure reduced the number of oocysts in a dose-dependent manner [13]. Together these early experiments all pointed in the same direction: that sulfadiazine exposure is detrimental for proper transmission of *Plasmodium* parasites in mosquitoes. However, the limitations of these experiments leave many important details unanswered.

Here we have investigated the effects of sulfadiazine treatment of mice and mosquitoes upon the transmission of *P. yoelii* gametocytes to *An. stephensi* mosquitoes. Specifically, we considered if the timing of exposure to sulfadiazine affects transmission, if pABA can help overcome treatment with sulfadiazine, and if treated parasites will mature *in vitro* as expected. We found that sulfadiazine exposure in the host or mosquito vector resulted in significantly decreased prevalence and intensity of infection in the mosquito midgut. Furthermore, we observed that providing excess pABA to mosquitoes in their sugar water only marginally rescued the effects of sulfadiazine exposure in the host. Finally, when sulfadiazine-treated parasites were cultured *in vitro* to produce ookinetes, no difference in the proportion of mature ookinetes was observed when sulfadiazine was washed out, indicating that the effects of sulfadiazine exposure may be reversible.

## Results

### Sulfadiazine treatment of the host limits transmission to mosquitoes

To first test if the standard treatment of infected mice with sulfadiazine for the selection of gametocytes had any effect on parasite transmission, mice were provided with standard drinking water or sulfadiazine-treated drinking water for two days leading up to the infectious blood meal to mosquitoes (schematic in Figure 1A). Following transmission to *An. stephensi* mosquitoes, we assessed the prevalence and intensity of infection under each condition seven days later. Parasites treated in the host with sulfadiazine were severely impacted in their ability to transmit to mosquitoes; in three of four biological replicates no transmission was observed when mosquitoes fed on sulfadiazine-treated mice (Figure 1B). In the fourth biological replicate (grey data points), mosquitoes that fed on sulfadiazine-treated mice were able to be infected, though at a significantly reduced intensity of infection, with significantly fewer oocysts per midgut observed (Figure 1C, Mann-Whitney unpaired t-test, p-value < 0.0001). There was no observed difference in the size or morphology of oocysts that did form in mosquitoes that fed on sulfadiazine treated mice (representative oocyst micrographs provided in Supplemental Figure 1).

**Figure 1.**
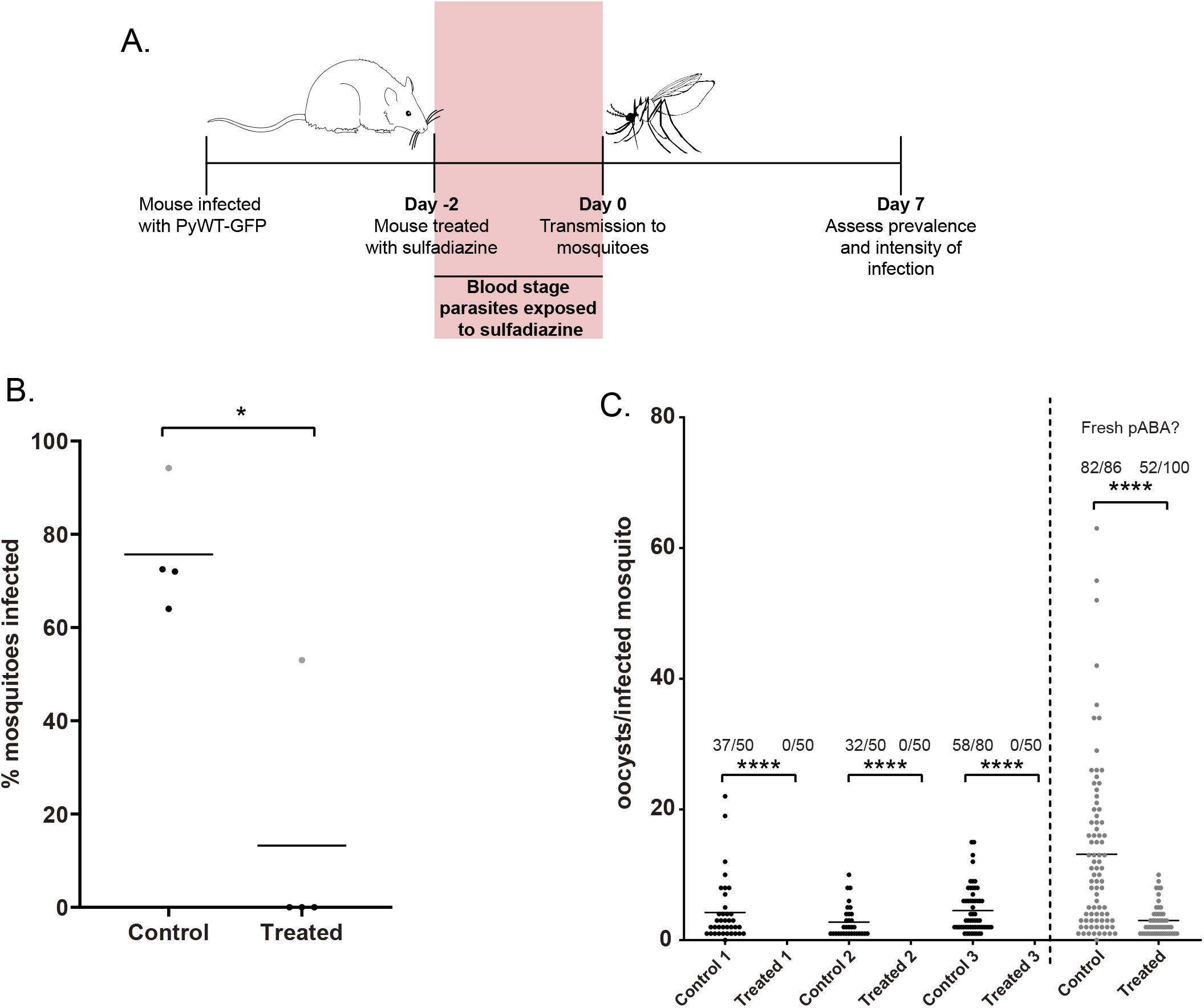
Sulfadiazine exposure in the host limits transmission to the mosquito vector. A. Mice infected with PyWT-GFP parasites were given standard drinking water (control) or water supplemented with 10 mg/L sulfadiazine (treated) for two days before mosquitoes were allowed to take an infectious blood meal. On day 7 post-blood meal, the percentage of mosquitoes infected (prevalence) (B) and the number of oocysts per infected mosquito (intensity of infection) (C) were assessed by live fluorescence microscopy. B. The average prevalence of infection for each biological replicate is represented by a data point, and the mean percentage of infected mosquitoes of all replicates is provided as a horizontal line. The grey data points correspond to the fourth replicate in Panel C. Mann-Whitney unpaired t-test was used for statistical analyses; * p-value < 0.05. C. The intensity of infection as measured by the number of counted oocysts per infected mosquito is plotted and the number of infected mosquitoes over the total number of mosquitoes counted for each sample is listed above each sample. Biological conditions for the final replicate (gray data points) may have been different, including supplementation of mosquitoes with fresher pABA (tested in Figure 2). Mann-Whitney unpaired t-test was used for statistical analyses; **** p-value < 0.0001.

### pABA supplementation of mosquitoes does not overcome exposure of parasites to sulfadiazine in the host

Because the fourth biological replicate (grey data points, Figure 1B-C) did result in limited transmission to mosquitoes, we considered whether there may have been different experimental conditions that allowed transmission of sulfadiazine-exposed parasites in this replicate. One such condition that may affect sulfadiazine treatment is the amount of pABA present in the mosquito vector, as sulfadiazine is a structural analog of pABA that acts as a competitive inhibitor of DHPS, and pABA supplementation of mosquitoes allows for increased numbers of oocysts to develop [13]. To determine if provision of fresh pABA to the mosquitoes could enable such transmission to occur, we replaced the pABA-supplemented sugar water daily using freshly dissolved pABA both before and after infectious blood meals taken from either control or sulfadiazine-treated mice. Consistent with this, we observed that fresh pABA supplementation enabled parasites to partially overcome exposure to sulfadiazine in the host and to successfully transmit to mosquitoes, albeit still at significantly lower levels than the untreated control (Figure 2B, Mann-Whitney unpaired t-test, p-value = 0.01). Moreover, the transmission intensity (oocysts per infected mosquito) was still significantly lower for mosquitoes that fed on treated mice than untreated mice (Figure 2C, Mann-Whitney t-test, p-value < 0.0001). Additionally, fresh pABA supplementation also improved the percentage of mosquitoes infected that fed on control mice as compared with routine pABA supplementation (compare Figure 1B and Figure 2B, average percent infected: 75.68 vs. 80.45). This indicates that the standard practice of sulfadiazine treatment of parasites in the rodent host to select for gametocytes is detrimental to parasite transmission, and that this practice may introduce unwanted artifacts that could complicate the study of these important transmission stage parasites.

**Figure 2.**
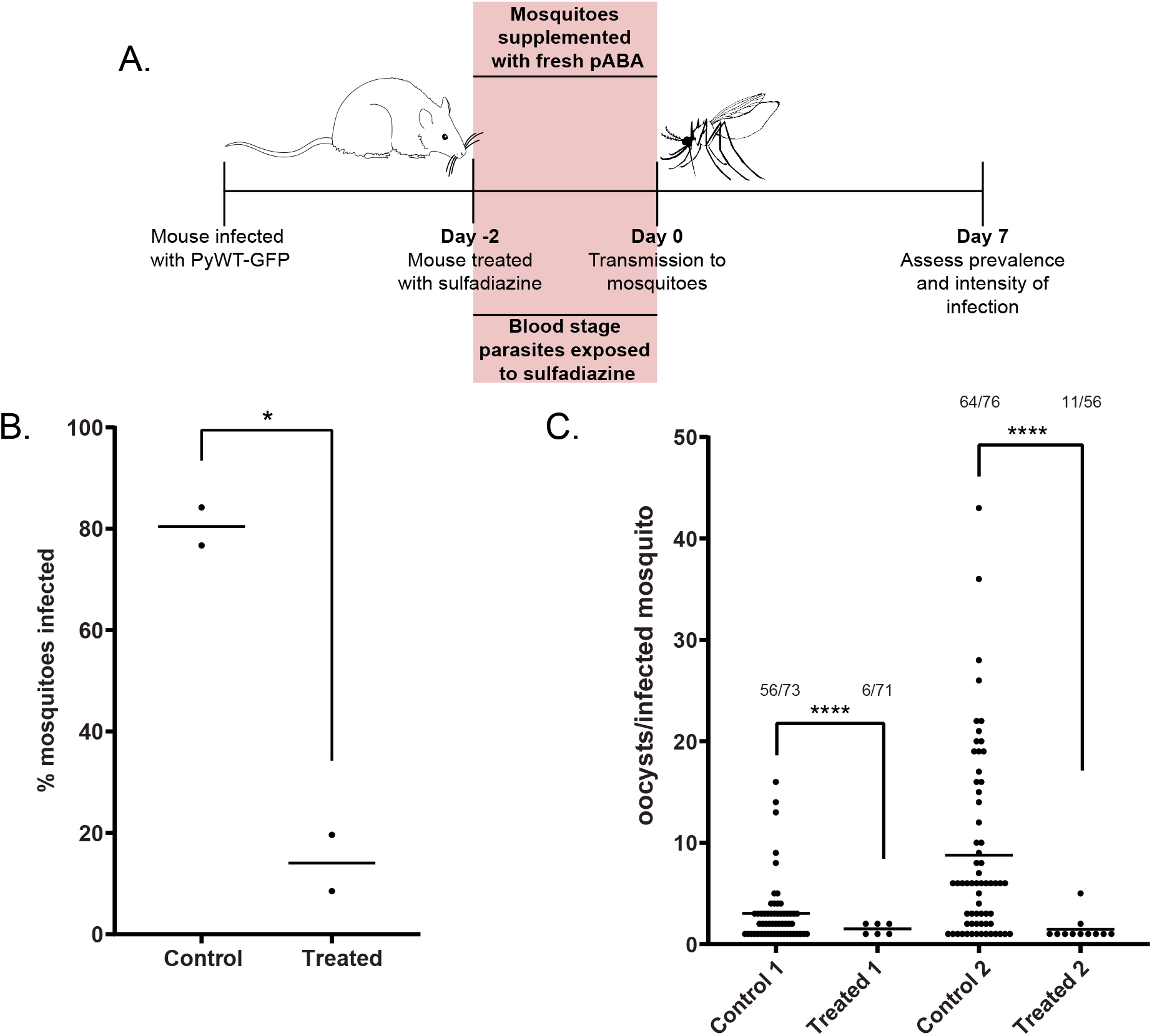
Fresh pABA supplementation can partially recover sulfadiazine exposure of parasites in the host. A. Mice infected with PyWT-GFP parasites were given standard drinking water (control), or water supplemented with 10 mg/L sulfadiazine (treated), two days before mosquitoes were allowed to take an infectious blood meal. Mosquito sugar water was supplemented daily with freshly diluted pABA (0.05% w/v) to test if fresh pABA can compete with sulfadiazine in the blood meal to recover parasite infection in the mosquito. On day seven post-blood meal, the percent of infected mosquitoes (prevalence) (B) and the number of oocysts per infected mosquito (intensity of infection) (C) were assessed by live fluorescence microscopy. B. The average prevalence of infection for each biological replicate is represented by a data point, and the mean percentage of infected mosquitoes of all replicates is provided as a horizontal line. Mann-Whitney unpaired t-test was used for statistical analyses; * p-value = 0.01. C. The intensity of infection, as measured by the number of counted oocysts per infected mosquito is plotted. The number of infected mosquitoes out of the total number of mosquitoes counted for each sample is listed above each sample. Mann-Whitney unpaired t-test was used for statistical analyses; **** p-value < 0.0001.

### Pre-treatment of mosquitoes with sulfadiazine reduces the intensity of mosquito infection

As sulfadiazine is bioavailable in the blood of the host [9], it will also be taken up along with the parasites during a blood meal. This would effectively extend the sulfadiazine exposure to the earliest mosquito stages as well. To test if parasites exposed only to sulfadiazine in the mosquito would have similar effects on transmission, we provided the mosquitoes with sulfadiazine in their sugar/pABA water for two days ahead of an infectious blood meal taken from untreated, infected mice (Figure 3A). We did not observe a statistically significant effect upon the prevalence of parasite transmission due to sulfadiazine exposure that was restricted to the mosquito midgut (Figure 3B, Mann-Whitney unpaired t-test, p-value = 0.400, not significant). Despite this, there was a significant reduction in the number of oocysts per infected mosquito observed when mosquitoes were treated with sulfadiazine (Figure 3C), indicating that sulfadiazine can affect the early mosquito stages (gametes, zygotes, ookinetes), as well as the sexual blood stage gametocytes.

**Figure 3.**
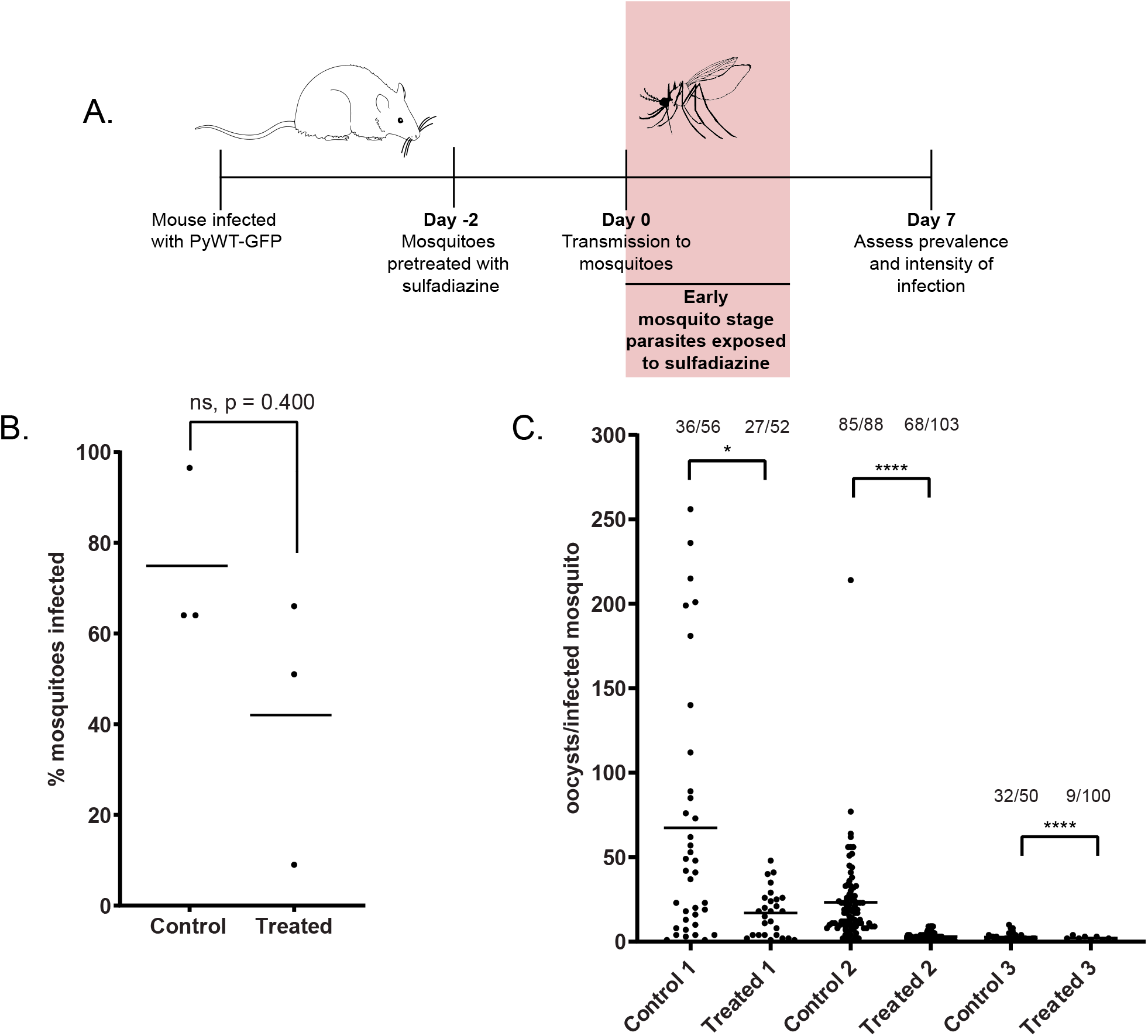
Sulfadiazine exposure to early mosquito stages decreases oocyst intensity. A. Mosquitoes were given standard sugar/pABA water, or sugar/pABA water supplemented with 10 mg/L sulfadiazine for two days leading up to an infectious blood meal from PyWT-GFP-infected mice, such that early mosquito stages are then exposed to sulfadiazine as they are taken up by the mosquito. On day 7 post-blood meal, the percentage of mosquitoes infected (prevalence) (B) and the number of oocysts per infected mosquito (intensity of infection) (C) were assessed by live fluorescence microscopy. B. The percentage of mosquitoes infected that fed on PyWT-GFP mice when mosquitoes were exposed to sulfadiazine (treated), or not (control) daily leading up to the infectious bloodmeal is plotted. The average percent infection for each biological replicate are represented by each data point, and the mean prevalence of all replicates is provided as a horizontal line. Mann-Whitney unpaired t-test was used for statistical analyses; ns = no significant difference, p-value > 0.05. C. The intensity of infection, as measured by the number of counted oocysts per infected mosquito is plotted. The number of infected mosquitoes out of the total number of mosquitoes counted for each sample is listed above each sample. Mann-Whitney unpaired t-test was used for statistical analyses; * p-value <0.05; **** p-value < 0.0001.

### *Sulfadiazine exposure in the host does not affect morphological development of* in vitro *ookinetes*

Because sulfadiazine treatment of the rodent host limited parasite development *in vivo* (Figures 1 and 2), and early mosquito stages were affected by sulfadiazine exposure in the mosquito midgut (Figure 3), we tested if sulfadiazine treatment had a reversible effect upon parasite development through these stages. To this end, mixed blood stage parasites from untreated or sulfadiazine-treated mice were collected using an Accudenz gradient, then resuspended and cultured *in vitro* in a defined medium containing pABA (1.0 mg/L; 7.299 μM) to assess fertilization and ookinete maturation (Figure 4A). Using differential interference contrast (DIC) microscopy, we did not observe any gross morphological differences in retorts or ookinetes that formed from either sulfadiazine-treated or untreated parasites (Figure 4B). Quantification of retort and ookinete stage parasites revealed no statistically significant differences in the proportions of retorts and ookinetes present in culture (Figure 4C, two-proportion Z-score test, p-value > 0.05, not significant).

**Figure 4.**
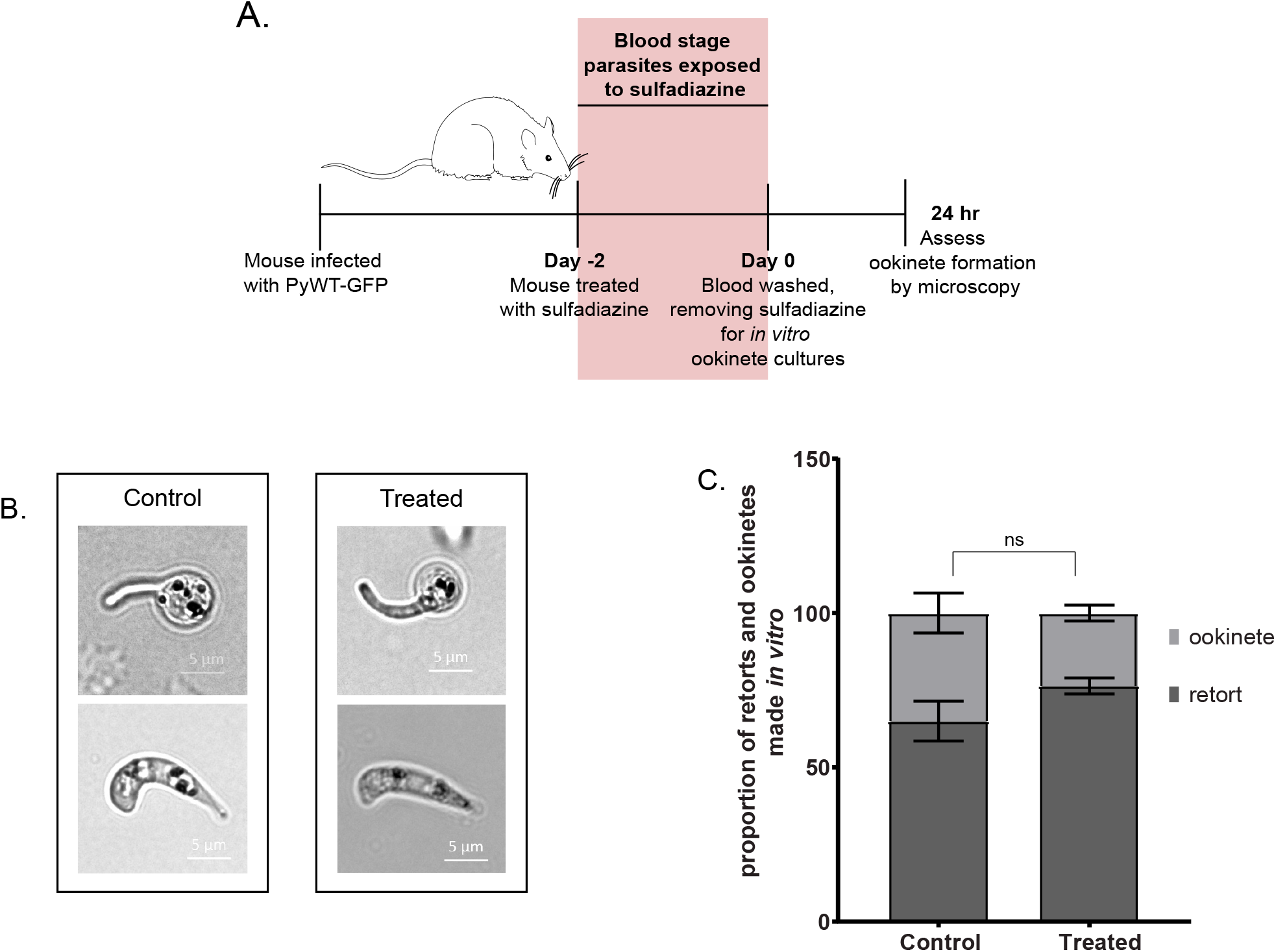
Sulfadiazine exposure to parasites in the host does not affect their ability to morphologically mature *in vitro*. A. Mice infected with PyWT-GFP parasites were given standard drinking water (control) or drinking water supplemented with 10 mg/L sulfadiazine (treated) for two days before parasite collection by exsanguination. Collected parasites were enriched by a discontinuous Accudenz gradient before being resuspended in sulfadiazine-free ookinete culture medium at room temperature. After 24 hours, the proportion of immature retorts and mature ookinetes was assessed by morphology. B. Representative images of retorts (top) or mature ookinetes (bottom) from control and treated parasites. C. The proportion of immature retorts and mature ookinetes observed in culture from blood of mice that was supplemented with sulfadiazine (treated) or not (control). A two proportion Z-score test was used for statistical analysis; ns = no significant difference, p-value > 0.05.

Taken together, these data demonstrate that treatment with sulfadiazine impacts not only asexual blood stage parasites, but also gametocytes and early mosquito stage parasites. This indicates that not only does the exposure of gametocytes to sulfadiazine affect transmission, but that exposure of the early mosquito stages to sulfadiazine within the mosquito midgut has a transmission blocking effect as well. Moreover, it is feasible that sulfadiazine can be introduced to the mosquito midgut via the rodent host or directly by the mosquito vector. Finally, consistent with sulfa drugs being competitive inhibitors of *Plasmodium* DHPS, we also conclude that the effects of sulfadiazine treatment upon *Plasmodium* development in early mosquito stage is reversible and can be at least partially overcome by competition by pABA supplementation of mosquitoes.

## Discussion

Sulfadiazine treatment is routinely used in rodent-infectious *Plasmodium* research labs to select for sexual blood stage gametocytes, the only life stage transmissible from host to vector. The enrichment of this stage away from other blood stage parasites is therefore critical to be able to robustly study the biology of host-to-vector transmission (epigenetic studies, transcriptomic studies, etc.). In particular, the development of novel transmission blocking strategies to prevent the further spread of malaria is dependent on a deep understanding of sexual stage biology. However, the underappreciated effect of gametocyte enrichment by sulfadiazine could be impacting these studies of parasite transmission.

Here we have shown that parasite transmission is impaired by sulfadiazine exposure of parasites in both the host and in the mosquito vector. The timing of exposure to sulfadiazine treatment is important and impactful, as mosquitoes that fed on sulfadiazine-treated mice likely have taken up sulfadiazine with their bloodmeal. In this scenario, if sulfadiazine was present in the blood bolus, it is feasible that early mosquito stage zygotes and ookinetes were exposed to the drug as well. Because sulfonamides are pABA analogs, we tested if treatment of mosquitoes with fresh pABA water could overcome sulfadiazine exposure. Ultimately, pABA supplementation of sulfadiazine-treated parasites resulted in only a partial restoration of transmission capability, as there were still significantly lower numbers of mosquitoes infected and lower numbers of oocysts per infected mosquito with sulfadiazine treated parasites, even upon fresh pABA supplementation. *In vitro*, sulfadiazine-exposed gametocytes can still develop into morphologically mature ookinetes. The blood used for *in vitro* culturing of ookinetes was enriched using an Accudenz gradient to collect infected red blood cells, so effectively any sulfadiazine in the whole blood collected from the mice was washed out. The excess pABA present in the ookinete culture media can then outcompete for binding to DHPS and allow the parasites to develop as expected.

It is possible that sulfadiazine treatment of the mouse or of mosquitoes before transmission may affect the mosquito midgut microbiome as well as the *Plasmodium* parasites. We did not observe adverse effects on mosquito survival with sulfadiazine supplementation. Though we have not directly studied the effects of sulfadiazine treatment on the mosquito midgut microbiome here, when placed in the context of previous studies, it is most plausible that the sulfadiazine-induced transmission defect we observed is parasite specific. Several studies that have explored these effects are worth noting. First, it was demonstrated that the mosquito midgut bacteria can have a negative effect on *Plasmodium* development in the mosquito, and that antibiotic treatment leads to higher parasite infection [21]. Additionally, it was shown that mosquitoes that fed on *Plasmodium-infected* blood containing penicillin and streptomycin had enhanced mosquito stage infections and reduced bacterial growth [22]. This antibiotic treatment could perhaps give the parasites a competitive advantage over the bacterial flora for nutrients in the mosquito midgut. If sulfadiazine treatment was adversely affecting the mosquito microbiome, we could similarly anticipate that the parasites would have enhanced development in the mosquitoes, rather than a defect such as what we have observed.

Recent work exposing mosquitoes to antimalarials, rather than antibiotics, may prove to be a new way to prevent new infections in natural transmission settings. For example, *Anopheles gambiae* exposed to atovaquone before an infectious bloodmeal results in a mosquito stage infection deficiency [23]. Testing more antimalarial drugs in this fashion could improve transmission blocking strategies for the elimination of malaria. This is another example of antimalarials taking on a new life after blood stage parasite resistance has emerged.

Finally, there are other means to enrich for gametocytes that may be preferable and less disruptive to transmission, like flow cytometry using gametocyte-specific antibodies or available fluorescent reporter lines using male- or female-enriched promoters [24–25], or magnetic enrichment [26–27]. Based upon these results, we strongly encourage their use over sulfadiazine selection to produce as minimally perturbed parasites as possible for the study of host-to-vector transmission.

## Materials and Methods

### Ethics Statement

All vertebrate animal care followed the Association for Assessment and Accreditation of Laboratory Animal Care (AAALAC) guidelines and was approved by the Pennsylvania State University Institutional Animal Care and Use Committee (IACUC# PRAMS201342678). All procedures involving vertebrate animals were conducted in strict accordance with the recommendations in the Guide for Care and Use of Laboratory Animals of the National Institutes of Health with approved Office for Laboratory Animal Welfare (OLAW) assurance.

### Use and Maintenance of Experimental Animals

Six-to-eight-week-old female Swiss Webster (SW) mice were used for all experiments in this work. *Anopheles stephensi* mosquitoes were reared and maintained at 24°C and 70% humidity under 12-hour light/dark cycles and were fed 0.05% w/v pABA-supplemented 10% (Sigma Aldrich, Cat#100536-250G) w/v sugar water. Mice were infected intraperitoneally with PyWT-GFP transgenic parasites that constitutively express GFP under a constitutive EF1 alpha promoter from the *pyp230p* dispensable locus (described previously,[28]).

### Treatment of Mice and Mosquitoes with Sulfadiazine

Mice were provided with standard drinking water before infection. After infection with PyWT-GFP parasites, the mice were kept on standard drinking water or were provided water supplemented with 10 mg/L sulfadiazine (VWR, Cat# AAA12370-30) for two days before the infectious blood meal to select for gametocytes. Mosquitoes were provided normal pABA/sugar water (0.05% w/v pABA, 10% w/v sugar) during rearing, and kept on normal pABA/sugar water, or supplied with pABA/sugar water supplemented with 10 mg/L sulfadiazine for two days before the infectious blood meal. After the blood meal, mosquitoes were again provided standard sugar/pABA drinking water for the duration of the *Plasmodium* infection.

### Host-to-Vector Transmission of *Plasmodium yoelii*

Mice infected with PyWT-GFP parasites were screened daily for parasitemia by Giemsa-stained thin blood smears and for the presence of male gametogenesis (visible as discrete exflagellation centers) via wet mount of a drop of blood incubated at room temperature for 8-10 minutes, as described previously [29]. On the peak day of exflagellation, the infected mice were anesthetized with a ketamine/xylazine cocktail and the mosquitoes were allowed to feed on the mice once for 15 minutes. Mosquito midguts were dissected seven days post-blood meal, and the prevalence and intensity of infection were assessed by differential interference contrast (DIC) and live fluorescence microscopy (Zeiss Axioscope A1 with 8-bit AxioCam ICc1 camera) using a 100X oil objective and processed by Zen 2012 (blue edition) imaging software.

### Production of *In Vitro* Ookinetes

Mice infected with PyWT-GFP parasites were supplied with standard drinking water or were treated with 10 mg/L sulfadiazine-treated drinking water to select for gametocytes for two days leading up to exsanguination. Parasitemia was assessed by Giemsa-stained thin blood smears and centers-of-movement were assessed to establish optimal experimental timing as described above. Blood was collected by cardiac puncture and then was maintained at 37°C in incomplete RPMI with 25mM HEPES and L-Glutamine (VWR Cat# 45000-412). Infected red blood cells were enriched by an Accudenz discontinuous gradient as previously described [30]. Ookinete cultures were generated as previously described, with modifications for *P. yoelii* [31–32]. Briefly, the Accudenz collected cells were added to *in vitro* ookinete media (RPMI 1640, 20% v/v FBS (Fisher Scientific, Cat#MT35011CV), 0.05% w/v hypoxanthine (Fisher Scientific, Cat#AC122010250), 100 μM xanthurenic acid (Sigma Aldrich, Cat# D120804-1G), pH 8.2 at 22°C) and were allowed to develop for 24 hours at room temperature. The pH was adjusted to pH 8.2 using KOH, rather than pH 7.5 for *P. berghei*, and cultures were maintained for 24 hours at ambient temperature (22°C), rather than a 19°C incubator. Retorts and ookinetes were then observed and quantified by DIC and fluorescence microscopy (Zeiss Axioscope A1 with 8-bit AxioCam ICc1 camera) using the 100X oil objective and processed by Zen 2012 (blue edition) imaging software.

### Statistical Analyses

Statistical differences in midgut oocysts infection numbers and numbers of infected mosquitoes were analyzed by Mann-Whitney unpaired t-test with p< 0.05 indicating statistical significance. P-values are listed where significant. The Mann-Whitney t-test was used because the data does not follow a normal distribution and control and experimental groups are independent of each other. A two proportion Z-score test was used for statistical analyses of ookinete and retort formation *in vitro* under no treatment conditions or with sulfadiazine treatment; p-value < 0.05 indicates significance. All statistical analyses were performed using Graphpad Prism (v8).

## Data Availability Statement

All data related to this study is provided in the manuscript and accompanying supplemental files.

## Acknowledgements

We thank Katarzyna Modrzynska (U. Glasgow) for helpful discussions related to the optimization of *in vitro* ookinete cultures for *P. yoelii*. We thank Mikaela Follmer for her work maintaining and rearing the Lindner laboratory mosquito colony. Finally, we thank members of the Lindner and Llinás laboratories for critical discussions of this work.

## Supplemental Figure Legend

**Supplemental Figure 1.**
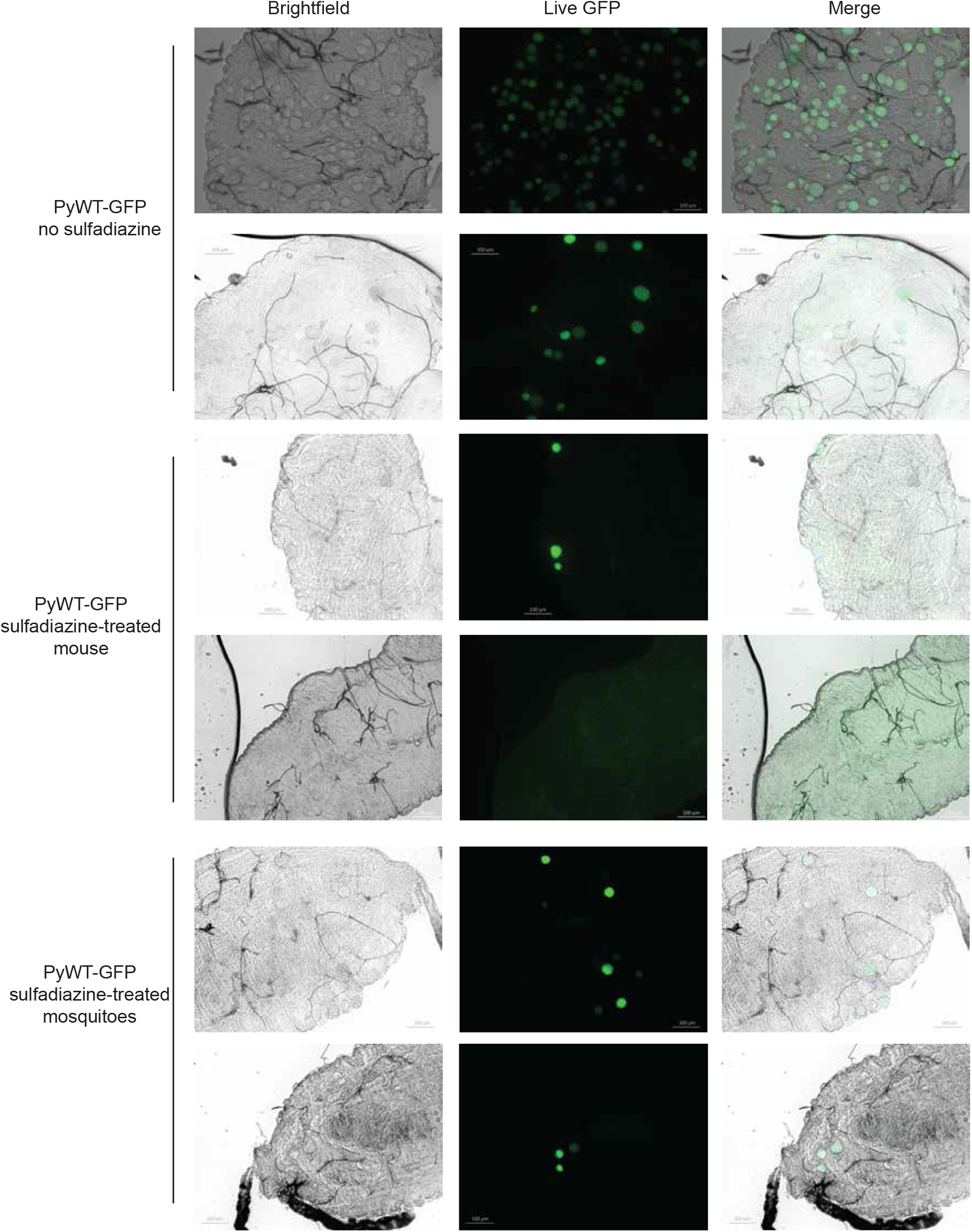
Representative images of midguts seven days after bloodmeal under control conditions, or with sulfadiazine supplementation of the rodent host or mosquito vector. Midgut infection with PyWT-GFP oocysts was assessed by live fluorescence microscopy seven days after the infectious bloodmeal. Scale bars are 100 μm.

